# Suppression of Inflammation Delays Hair Cell Regeneration and Functional Recovery Following Lateral Line Damage in Zebrafish Larvae

**DOI:** 10.1101/2020.02.26.962753

**Authors:** Ru Zhang, Xiao-Peng Liu, Ya-Juan Li, Ming Wang, Lin Chen, Bing Hu

**Author notes:** Corresponding authors. Bing Hu, PhD, Lab of Neurodevelopment and Repair, University of Science and Technology of China, Hefei 230027, China., Lin Chen, PhD, Auditory Research Laboratory, University of Science and Technology of China, Hefei 230027, China.

## Abstract

**Background:** Human cochlear hair cells cannot spontaneously regenerate after loss. In contrast, those in fish and amphibians have a remarkable ability to regenerate after damaged. Previous studies focus on signaling mechanisms of hair cell regeneration, such as Wnt and Notch signals but seldom on the fact that the beginning of regeneration is accompanied by a large number of inflammatory responses. The detailed role of this inflammation in hair cell regeneration is still unknown. In addition, there is no appropriate behavioral method to quantitatively evaluate the functional recovery of lateral line hair cells after regeneration.

**Results:** In this study, we found that when inflammation was suppressed, the regeneration of lateral line hair cells and the recovery of the rheotaxis of the larvae were significantly delayed. Calcium imaging showed that the function of the neuromasts in the inflammation-inhibited group was weaker than that in the non-inflammation-inhibited group at the Early Stage of regeneration, and returned to normal at the Late Stage. Calcium imaging also revealed the cause of the mismatch between the function and quantity during regeneration.

**Conclusions:** Our results, meanwhile, suggest that suppressing inflammation delays hair cell regeneration and functional recovery when hair cells are damaged. This study may provide a new knowledge for how to promote hair cell regeneration and functional recovery in adult mammals.

## Background

Deafness and hearing defects are usually caused by loss of sensory hair cells or defect of auditory function. The loss of hair cells is result of aging, infection, genetic factors, hypoxia, autoimmune disorder, ototoxic drugs or noise exposure. Unfortunately, including humans, hair cells cannot regenerate in mammals (Oesterle and Stone, 2008; Yorgason. et al., 2006). In contrast, hair cells in some non-mammalian vertebrates have a remarkable ability to regenerate, such as birds, reptiles, amphibians and fish (Matsui. and Cotanchea., 2004; Popper and Hoxter, 1984; Stone. and Rubel., 2000). It could suggest that if we figure out the mechanism of hair cell regeneration in these species, we probably can promote hair cell regeneration in mammals.

When hair cells are damaged, support cells proliferate into both hair cells and support cells, or convert into hair cells directly (Baird et al., 1996; Lopez-Schier and Hudspeth, 2006; Raphael, 1992; Roberson et al., 2004). Hair cell regeneration is finely regulated by the interaction of multiple signaling pathways, such as Notch signaling (Ma et al., 2008; Mizutari et al., 2013), Wnt/b-catenin signaling (Aman and Piotrowski, 2008; Chai et al., 2012; Shimizu et al., 2012), Fgf signaling (Aman and Piotrowski, 2008; Nechiporuk and Raible, 2008), retinoic acid (Rubbini et al., 2015) and so on. In the process of hair cell damaged, it is accompanied by a lot of inflammatory reaction, which has been found to play a role in tissue regeneration in recent years (Mescher, 2017). For example, macrophages are considered having main function in the inflammatory resolution stage and being required for fin regeneration (Li et al., 2012) and hair cell regeneration in zebrafish (Carrillo et al., 2016). In addition, it has been confirmed that neutrophils in mice play a central role in inflammation-induced optic nerve regeneration (Kurimoto et al., 2013).

In recent years, zebrafish (Danio rerio) has become an ideal model for studying inflammation and hair cell regeneration because it has conservative innate immunity (Renshaw and Trede, 2012) and strong regeneration ability in lateral line system (Lush and Piotrowski, 2014) which makes zebrafish larvae to perceive the change of surrounding flow, detect their prey and avoid predators (Coombs. et al., 2014; Dijkgraaf, 1962). The lateral system of a larva is composed of neuromasts which located on the surface of the body. The neuromasts on the head consist of the anterior lateral line system (aLL) and the ones along the body comprise the posterior lateral line system (pLL)(Thomas et al., 2015). The center of the neuromast is composed of hair cells and they are surrounded by support cells and mantle cells. At the top of the hair cells, rows of short stereocilia and a long kinocilium extend out of the body called the hair bundle and are covered in a gelatinous cupula. The arrangement of stereocilia and kinocilium determines the polarity of hair cells and the polarity of the hair cells is planar cell polarity (PCP), which is arranged symmetrically (Flock and Wersall, 1962), half in each direction.

When hair bundles are deflected, hair cells release transmitters and cause exciting spikes in afferent neurons (Dijkgraaf, 1962). And then, larvae show a robust behavior called rheotaxis (Olszewski et al., 2012). This behavior can be applied to evaluate the function of hair cells (Suli et al., 2012).

In recent years, calcium imaging has become a popular method to measure the function of neural cells in detail and quantitatively (Zhang et al., 2016). When the mechanical hair bundle deflected, calcium and other cations enter into cytoplasm through mechanotransduction channels. It changes the membrane potential and activates voltage-gated calcium channels which allow rapid calcium inflow to trigger synaptic transmission. GCaMPs, a genetically-encoded calcium indicator(GECIs), are single fluorescent proteins, which can bind calcium directly and alter conformation to respond the change of calcium concentration (Tian et al., 2012). These significant, activity-dependent signals can reflect the function of hair cells in a single neuromast (Zhang et al., 2018; Zhang et al., 2016).

Previous research has found that the deletion of macrophages by morpholino leads to the delay of hair cell regeneration (Carrillo et al., 2016). However, does it still cause the delay of hair cell regeneration when the macrophages are intact, and the pro-inflammatory factors are suppressed as the hair cells are damaged? Is there any delay in the functional recovery of the lateral line?

In order to figure out the above problems, we used an anti-inflammatory agent, BRS-28, to suppress the inflammation when hair cells are damaged by copper. BRS-28 is a derivative of 5α-cholestan-6-one, which was confirmed to be a remarkably suppressor of the production of pro-inflammatory factors, such as NO, TNF-α, IL-1β, iNOS and cox-2 (Yang et al., 2014). We count the number of neutrophils and macrophages in Tg(corola-eGFP; lyz-Dsred) transgenic line. Then, AB/WT zebrafish larvae were used to count the number of regenerated hair cells. Since there is no appropriate behavioral method to quantitatively evaluate the function of lateral line hair cells, we designed and built devices to test rheotaxis behavior in AB/WT larvae. A behavioral analysis software was applied for quantitative evaluation of rheotaxis, so as to reflect the holistic functional recovery of the posterior lateral line. Finally, the function of the regenerated hair cells in a single neuromast was evaluated by the method of calcium imaging in Huc:h2b-gcamp6f transgenic line.

## Results

### CuSO_4_ damaged hair cells in lateral line of zebrafish

Sensory hair cells in a 6-day post fertilization (dpf) AB/WT zebrafish larva were labeled with 0.05% DASPEI clearly (**Fig. 1A**). L2, LII3, L3 neuromasts (circles in **Fig.1 A**) were three of the posterior lateral neuromasts, which located along the flat truck body and easily to be observed. A lateral view of the neuromasts showed the elongated kinocilia extending from the body (**Fig. 1B**). The neuromasts are consisted of hair cells surrounded by support cells, which are surrounded by mantle cells (**Fig. 1C**). In order to study the effects of inflammation on hair cell regeneration, we established a hair-cell-damaged model. Hair cells were damaged completely, when treated with 5 μM CuSO_4_ for 1 h (**Fig. 1D**). Labeled with 0.05% DASPEI, hair cells displayed close arrangement and clear boundary. Only treated with CuSO_4_ solution for 20 min, hair cells became loose and unclear which suggested that they were already injured. The number of hair cells decreased with weaker fluorescence intensity and obscure cell boundary at 40 min. Hair cells were completely disappeared at 60 min, indicating that they had been completely damaged. TUNEL assay revealed the missing hair cells underwent apoptosis (**Supplementary Fig. 1**). After being transferred to embryo medium (EM), the number of hair cells quickly returned to normal (**Fig. 1E**).

**Fig. 1.**
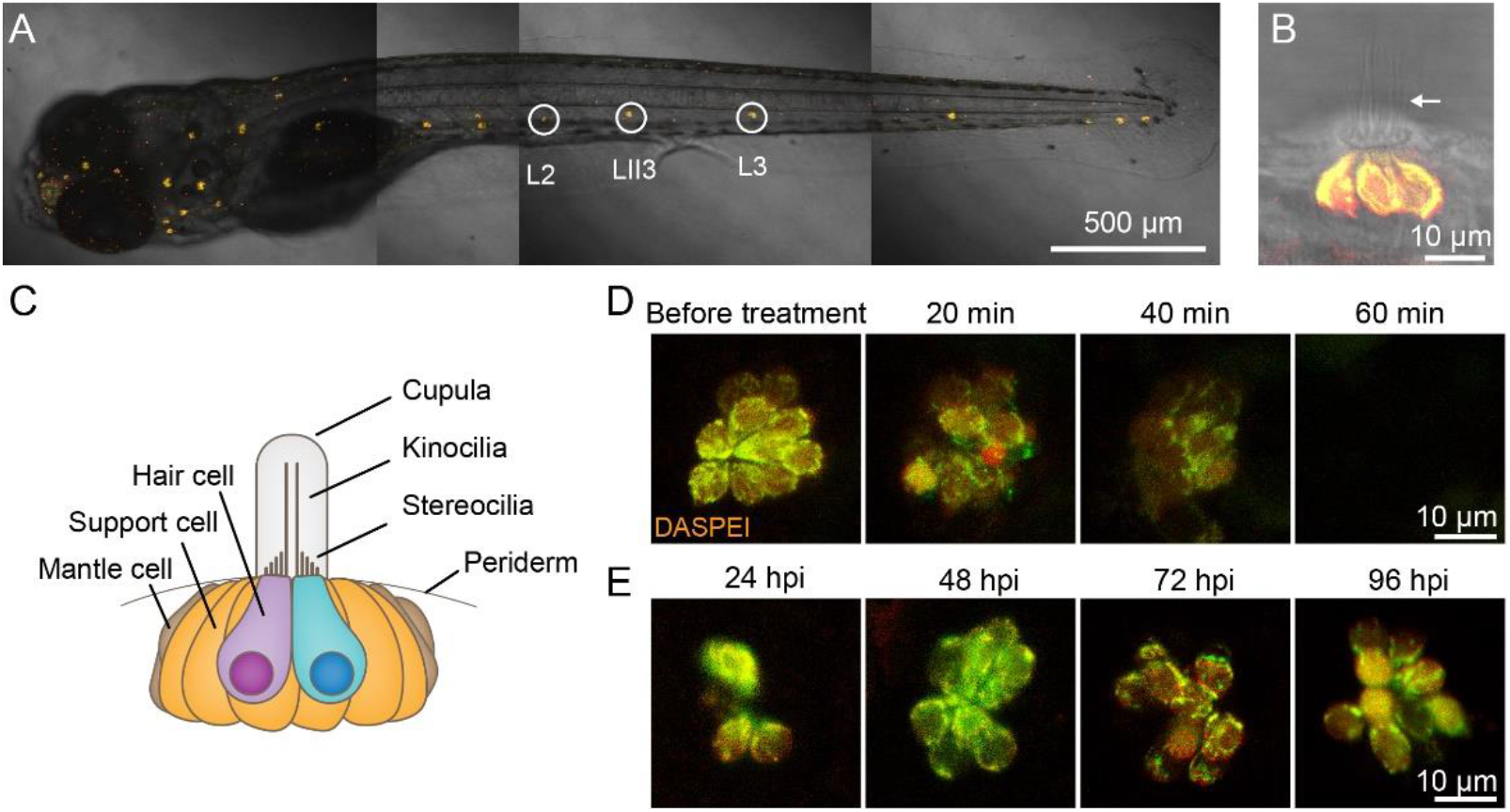
CuSO_4_ damaged hair cells in lateral line of zebrafish. **(A)** Lateral line hair cells in a 6 day post fertilization (dpf) AB/WT zebrafish larvae are labeled with 0.05% DASPEI. L2, LII3 and L3 neuromasts are marked with circles. Scale bar represents 500 μm. **(B)** Lateral view of a neuromast shows sensory hair cells in the center labeled with DASPEI and a bundle of kinocilia (arrow) extend out of the periderm. Scale bar represents 50 μm. **(C)** A cartoon illustrating the structure of the neuromast. **(D)** Time lapse imaging shows that when merged in 5 μM CuSO_4_ solution, hair cells were gradually injured and damaged within 60 min. Scale bar represents 10 μm. **(E)** DASPEI staining displays that hair cells regenerate completely within 96 hours post injured (hpi). Scale bar represents 10 μm.

### BRS-28 reduced the number of neutrophils and macrophages migrating to the injured neuromasts

Neutrophils (**Fig. 2B, C**, blue arrows) and macrophages (**Fig. 2B, C**, white arrows) could be marked and distinguished in larvae of Tg(corola-eGFP; lyz-Dsred) transgenic line (**Supplementary Fig. 2**). Normally, neutrophils and macrophages were almost absent from the neuromasts (example, **Fig. 2A**). When treated with CuSO_4_ solution, hair cells were damaged. Neutrophils and macrophages migrated to the neuromasts within 1 hours (example, **Fig. 2B**). When larvae were immerged in BRS-28, an anti-inflammatory agent, before treated with CuSO_4_ solution, less neutrophils and macrophages migrated to the damaged neuromasts (example, **Fig. 2C**). When the inflammation suppressed, the numbers of neutrophils appeared around the damaged neuromasts were lower at 0.5,1,3 and 4 h after adding the CuSO_4_ solution in BRS+CuSO_4_ group than in CuSO_4_ group (**Fig. 2D**). In addition, we observed BRS+CuSO_4_ group had fewer macrophages at 0.5, 1, 2 and 3 h than CuSO_4_ group (**Fig. 2E**). Collectively, the data strongly suggested that BRS-28 reduced the number of neutrophils and macrophages migrating to the injured neuromasts. It was worth noting that compared with control, there was no significant difference in the numbers of neutrophils and macrophages between CuSO_4_ group and BRS+CuSO_4_ group at 5 and 6 h, indicating that the inflammation was almost resolved.

**Fig. 2.**
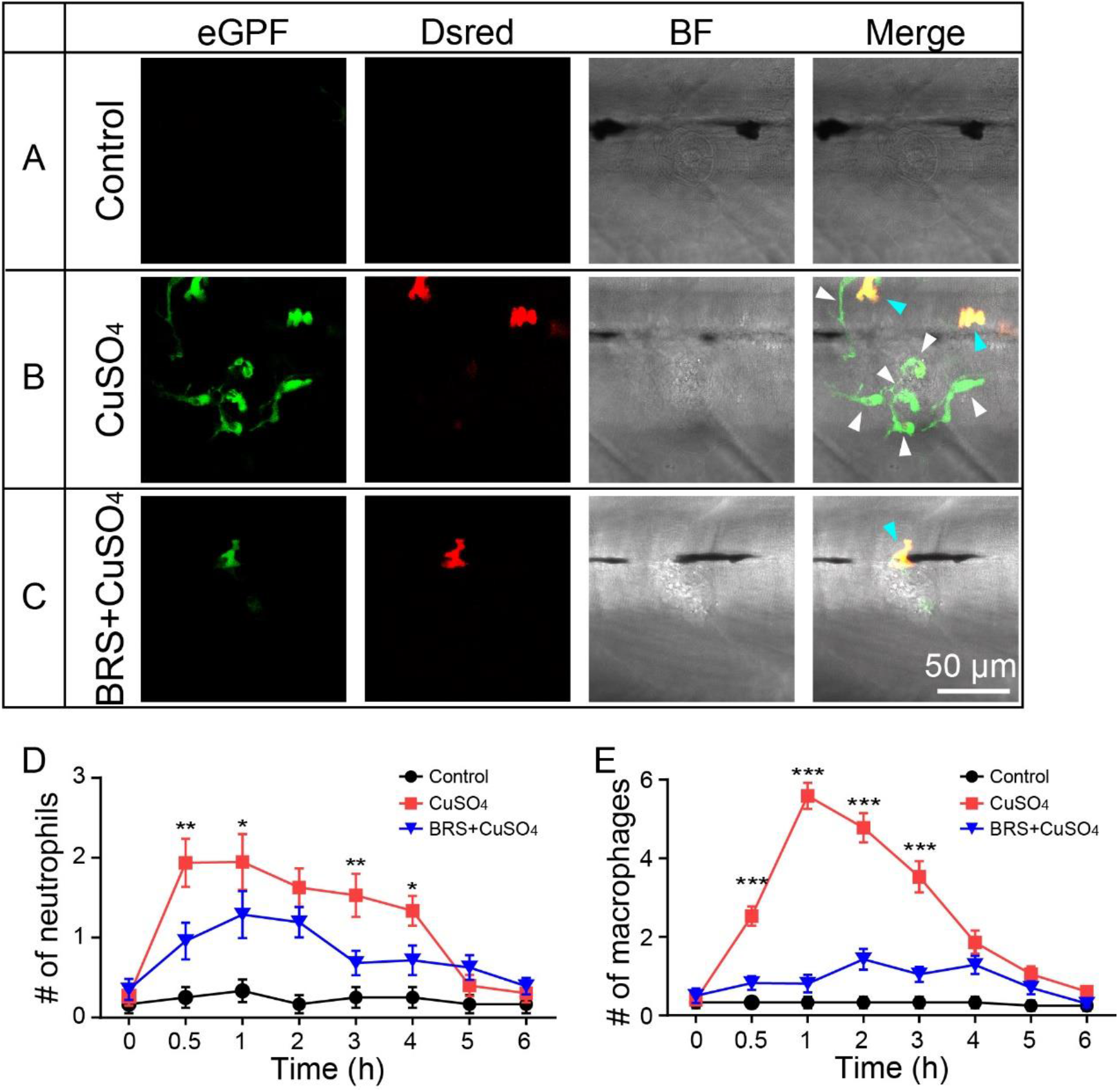
BRS-28 reduces the number of neutrophils and macrophages migrating to the injured neuromasts. **(A-C)** Live imaging (×40) displays the regions of L3 neuromasts of larvae at GFP channel, Dsred channel, and bright field (BF) channel and superimposed image in different group. Neutrophils (show both green and yollow fluorescence, indicated by white arrows) and macrophages (show olny green fluorescence, indicated by blue arrows) around the neuromasts can be observed in Tg(corola-eGFP; lyz-Dsred) larvae. They are almost absent from the neuromasts in Control group **(A)**. Many neutrophils and macrophages migrate to injured neuromasts in CuSO_4_ group **(B)** while fewer neutrophils and macrophages migrate to injured neuromasts in BRS+CuSO_4_ group **(C)**. The image is captured after adding CuSO_4_ solution for 1 h. Scale bar represents 50 μm. **(D-E)** Line charts reveal decreased numbers of neutrophils **(D)** and macrophages **(E)** within a radius of 50 μm from the center of neuromasts at different time points after adding CuSO_4_ in BRS+CuSO_4_ group (n≥16) than CuSO_4_ group (n≥15). Control group (n≥11) is observed at the same time points. To **(D)** and **(E)**, comparisons were performed by using two-way ANOVA, with Tukey’s multiple comparisons test. All Error bars show mean ± S.E.M., *** P < 0.001, **P < 0.01,*P < 0.05.

### Suppressing inflammation delayed hair cell regeneration

In order to investigate whether the regeneration of hair cells were delayed after suppressing inflammation, we observed hair cells in the L2, LII3 and L3 neuromasts. We found that the regeneration of hair cells was delayed after the inflammation was suppressed by the inflammatory inhibitor, BRS-28. Live imaging showed regenerated hair cells in CuSO_4_, BRS+CuSO_4_ group at 24, 48 and 96 hours post injured (hpi) by CuSO_4_(**Fig. 3A**). Control group was showed at the same time point. Further analysis revealed that the numbers of regenerated hair cells were significantly decreased in BRS+CuSO_4_ group than that in CuSO_4_ group at 16 hpi (P=0.0061), 24 hpi (P=0.0021) and 48 hpi (P<0.0001) (**Fig. 3B**, n = 30 neuromasts). These results indicated that the regeneration of hair cells was delayed in BRS+CuSO_4_ group within 48 hpi. Compared with Control group, there was no difference in the number of hair cells between CuSO_4_ group and BRS+CuSO_4_ group at 96 hpi, suggesting that hair cells were regenerated to the normal level at 96 hpi. We also analyzed the number of hair cells when only treated with BRS-28 (BRS group) without hair cell damage. As expected, BRS group had no difference compared with Control group at any time point, excluding the effect of BRS-28 on hair cells.

**Fig. 3.**
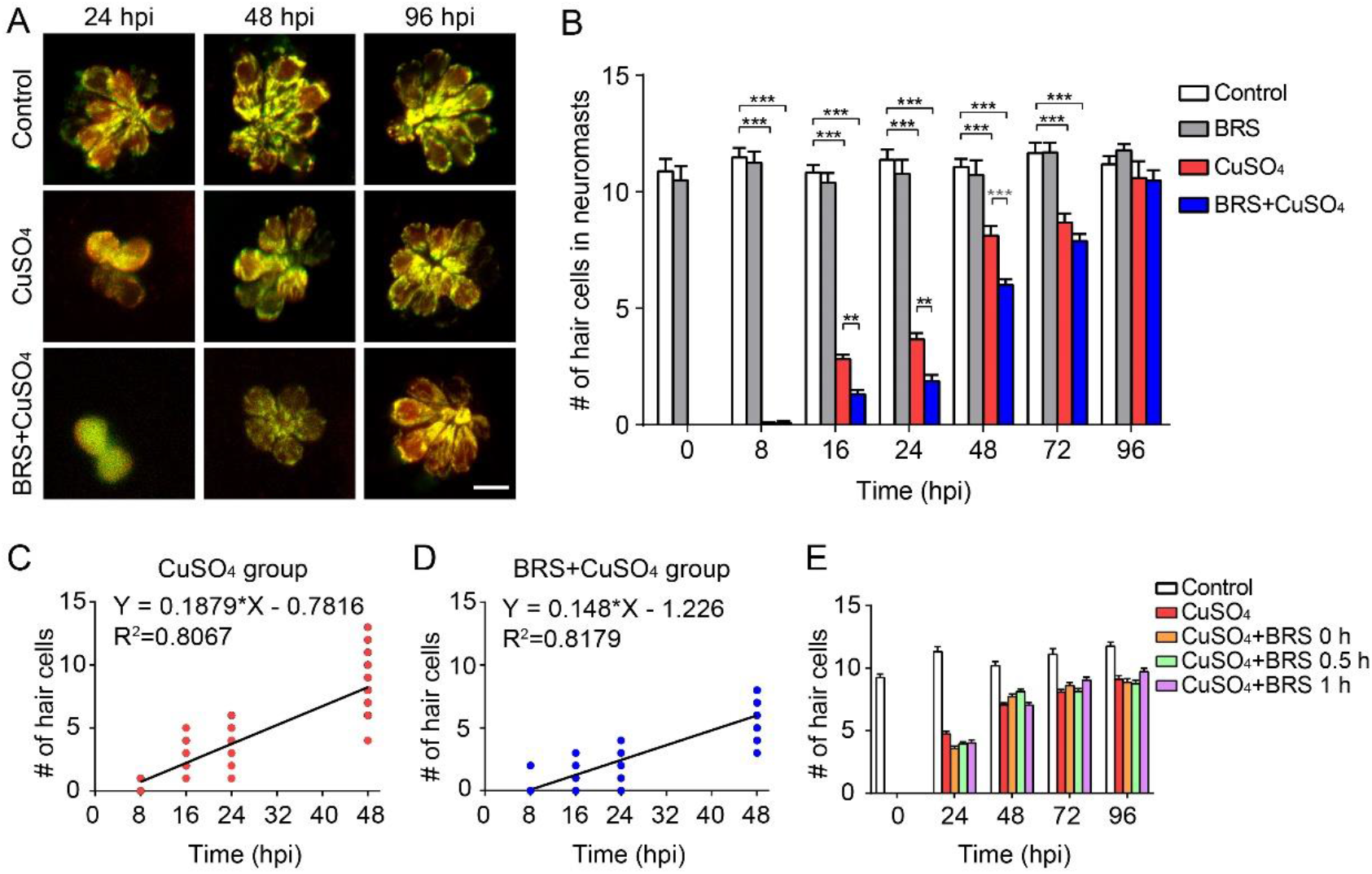
Suppressing inflammation delays hair cell regeneration. **(A)** Real-time imaging (×40) displays regenerated hair cells in the CuSO_4_ and BRS+CuSO_4_ group at 24, 48 and 96 hpi. Control group is taken at the same time point. Scale bar represents 10 μm. **(B)** The numbers of regenerated hair cells were significantly decreased in BRS+CuSO_4_ group than that in CuSO_4_ group at 16 (P=0.0061), 24 (P=0.0021) and 48(P<0.0001) hpi. At 96 hpi, hair cells in both CuSO_4_ group and BRS+CuSO_4_ group regenerated to normal levels. Linear analysis in CuSO_4_ group **(C)** and BRS+CuSO_4_ group **(D)** were conducted on the number of regeneration within 48 hours. The slope in CuSO_4_ group (0.1879) is higher than that in BRS+CuSO_4_ group (0.148) and x-intercept in CuSO_4_ group (4.16) is higher than that in BRS+CuSO4 group (8.287). **(E)** When delay the time window of inflammatory suppression, there is no delay in the regeneration of hair cells. BRS-28 was added at the same time as CuSO_4_ (CuSO_4_+BRS 0 h group), or 30 minutes after the addition of CuSO_4_ (CuSO_4_+BRS 0.5 h group), or 1 hour after the addition of CuSO_4_ (CuSO_4_+BRS 1 h group)(n≥27 neuromasts in each time point of each group). To **(B)** and **(E)**, comparisons were performed by using two-way ANOVA, with Tukey’s multiple comparisons test. All Error bars show mean ± S.E.M., *** P < 0.001, **P < 0.01,*P < 0.05.

Since hair cells did not regenerate at a uniform rate, we defined the time of regeneration into two periods: the Early Stage which includes the time from 0 to 48 hpi and the Late Stage which includes the time after 48 hpi. The regeneration of hair cells was fast in the Early Stage and slow in the Late Stage. Linear analysis was conducted on the number of hair cell regeneration in the Early Stage. The slope in CuSO_4_ group (0.1879) was higher than that in BRS+CuSO_4_ group (0.148), meanwhile, x-intercept in CuSO_4_ group (4.16) was higher than that in BRS+CuSO_4_ group (8.287) (**Fig. 3C, D**). These implied that the hair cell regeneration in BRS+CuSO_4_ group may begin later and slower than that in CuSO_4_ group.

To explore whether the time window of inflammatory suppression had contribute to delayed regeneration, we changed the start time of BRS-28 treatment. We found that compared with the CuSO_4_ group, whether BRS-28 was added at the same time as CuSO_4_ (CuSO_4_+BRS 0 h group), or 30 minutes after the addition of CuSO_4_ (CuSO_4_+BRS 0.5 h group), or 1 hour after the addition of CuSO_4_ (CuSO_4_+BRS 1 h group) (**Fig. 3E**), there was no statistical difference on the number of regenerated hair cells.

To sum up, the regeneration of hair cells in lateral line was delayed after the inflammation was suppressed by the inflammatory inhibitor BRS-28.

### The functional recovery of the lateral line system was delayed when inflammation was suppressed

Since the rheotaxis could be a suitable functional readout of the lateral line, we designed a behavioral device to test the rheotaxis of zebrafish (**Fig. 4A**, see details in Materials and Methods). Larvae were placed from the right platform, and they sense the water flow direction from the right to the left. **Figure 4B, C** were two examples of the larval rheotaxis processed by behavioral analysis software: the former was a larva with excellent rheotaxis (**Fig. 4B**) while the latter was a larva performed failure in the rheotaxis test (**Fig. 4C**). The left panels in these two examples showed the swimming track of this larva. The behavioral analysis software mapped its movement path of larvae by line segment. The color of the line segment represented the direction of movement of the larvae. All the movements from right to left were represented by purplish or red segments, where purple indicated that the velocity along the flow direction was greater than or equal to the flow velocity, and red indicated that the velocity along the flow direction was less than the flow velocity. All the movements from left to right were represented by green segments, and the higher the brightness was, the faster the speed was. The right panels displayed the motion vector. The lengths of the blue segments represented the distance of each movement, and the direction of the blue segment represented the direction of that movement. The length of the red line segment was the ratio of motion vectors sum to the motion arithmetic sum and the direction was the direction of the sum of the vectors.

**Fig.4.**
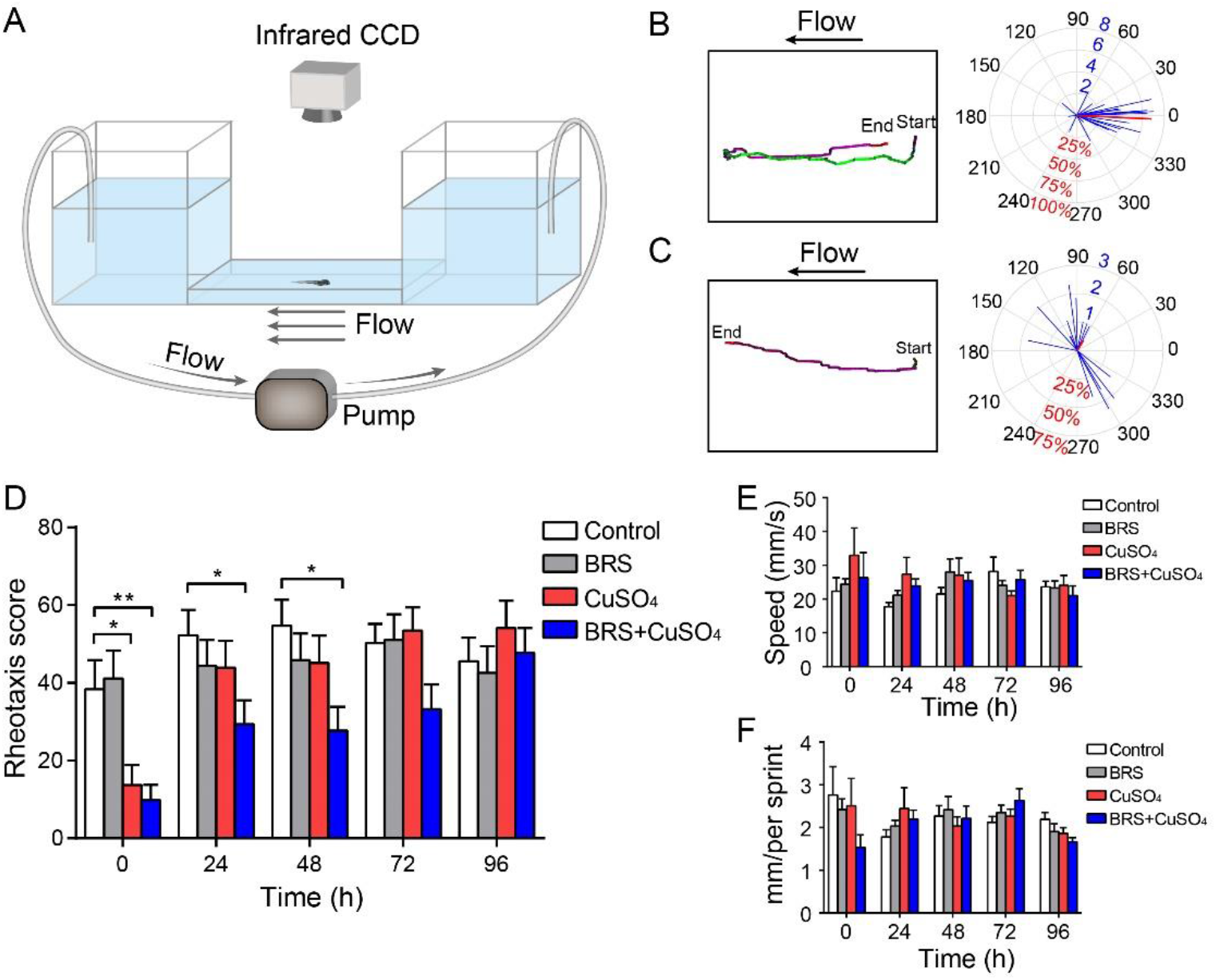
The recovery of the functional of lateral line system was delayed when inflammation was suppressed. **(A**) A U-shaped tank was designed to test the rheotaxis behavior of larvae. A peristaltic pump was used to form flow at the bottom of the tank. Larvae were placed from the right platform, and they sense the water flow from right to left. Rheotaxis perform was recorded by an infrared CCD. A larva with excellent rheotaxis **(B)** and a larva with poor rheotaxis **(C)** were analyzed by behavioral analysis software. Moving traces were plotted in left panels and the motion vector were displayed in right panels. The lengths of the blue segments represented the distance of each movement, and the direction of the blue segment represented the direction of that movement. The length of the red line segment was the ratio of motion vectors sum to the motion arithmetic sum and the direction was the direction of the sum of the vectors. **(D)** Rheotaxis score revealed that at 24 and 48 hpi, the rheotaxis of BRS+CuSO_4_ group was significantly lower than that of Control group. On the contrary, the rheotaxis of CuSO_4_ group was not significantly different from that of Control group within 24 hpi. The speed **(E)** and distance **(F)** of larvae swimming at each time were consistent within different times and between different groups. To **(D-F)**, comparisons were performed by using two-way ANOVA, with Tukey’s multiple comparisons test. All Error bars show mean ± S.E.M., **P < 0.01,*P < 0.05.

When the red segment was long and had a small angle of 0 degree, it indicated that the motion of the larva was consistent with the opposite direction of flow. It represented that the larva had a good rheotaxis, indicating its lateral line system executed its function very well. Therefore, the software reported a high score. On the contrary, when the red segment was short or had a small angle of 180 degree, it indicated that the larva moved randomly and had a poor rheotaxis, indicating its lateral line system had poor function. In this case, the software reported a low score. The scores reported by the software were plotted into bar charts and showed in **Figure 4D**. After the hair cells were damaged by CuSO_4_, there was poor rheotaxis in both CuSO_4_ group and BRS+CuSO_4_ group. At 24 and 48 hpi, the rheotaxis of BRS+CuSO_4_ group was significantly lower than that of Control group. On the contrary, the rheotaxis of CuSO_4_ group was not significantly different from that of Control group within 24 hpi. Therefore, it indicated that the functional recovery of lateral line system was delayed in BRS+CuSO_4_ group. The rheotaxis of BRS group at each time point was not different from that of Control group, suggesting that BRS-28 alone had no significant effect on the rheotaxis. In addition, we noted that the speed and distance of each movements were consistent within different times and between different groups: both were stable at around 22 mm/s (**Fig. 4E, F**), which indicated that BRS-28 or CuSO_4_ did not affect the movement of zebrafish.

We concluded that the regenerated hair cells still had the ability to sense water flow, but the functional recovery of lateral line system was delayed when inflammation was suppressed.

### Calcium imaging revealed the function of a single neuromast after hair cell regeneration

Since we found a mismatch between the function of the lateral line and the amount of hair cell regeneration, that is, after the zebrafish lateral line was damaged by copper sulfate, it took 96 h for the hair cells to return to normal, while the flow ability returned to normal at 24 h. The function of a single neuromast can be evaluated by observing its calcium activity (Zhang et al., 2016). The L3 neuromast, located in flat trunk, was stimulated by water flow from an electrode (**Fig. 5A**). Since hair cells had polarities, the yellow and green hair cells represented opposite polarities. Chou et al. reported that the polarity of the L3 neuromast is parallel to the anterior-posterior body axis (Chou et al., 2017). Thus, by adjusting the direction of the electrode, water was controlled to flow in two directions: anterior to posterior (A-P) direction or posterior to anterior (P-A) direction. We found that not all hair cells responded to the water flow, and only some hair cells were active (example, **Fig. 5B**, circled cells). These active cells only responded to stimulus in one direction: P-A direction (**Fig. 5C**, yellow ones and yellow circles in **Fig. 5B**) or A–P direction (**Fig. 5D**, green ones and green circles in **Fig. 5B**). Because the neuromasts were stereoscopic, some of the active hair cells were far from this focal plane (dashed circles in **Fig. 5B**) and were not included in subsequent fluorescence intensity analysis.

**Fig.5.**
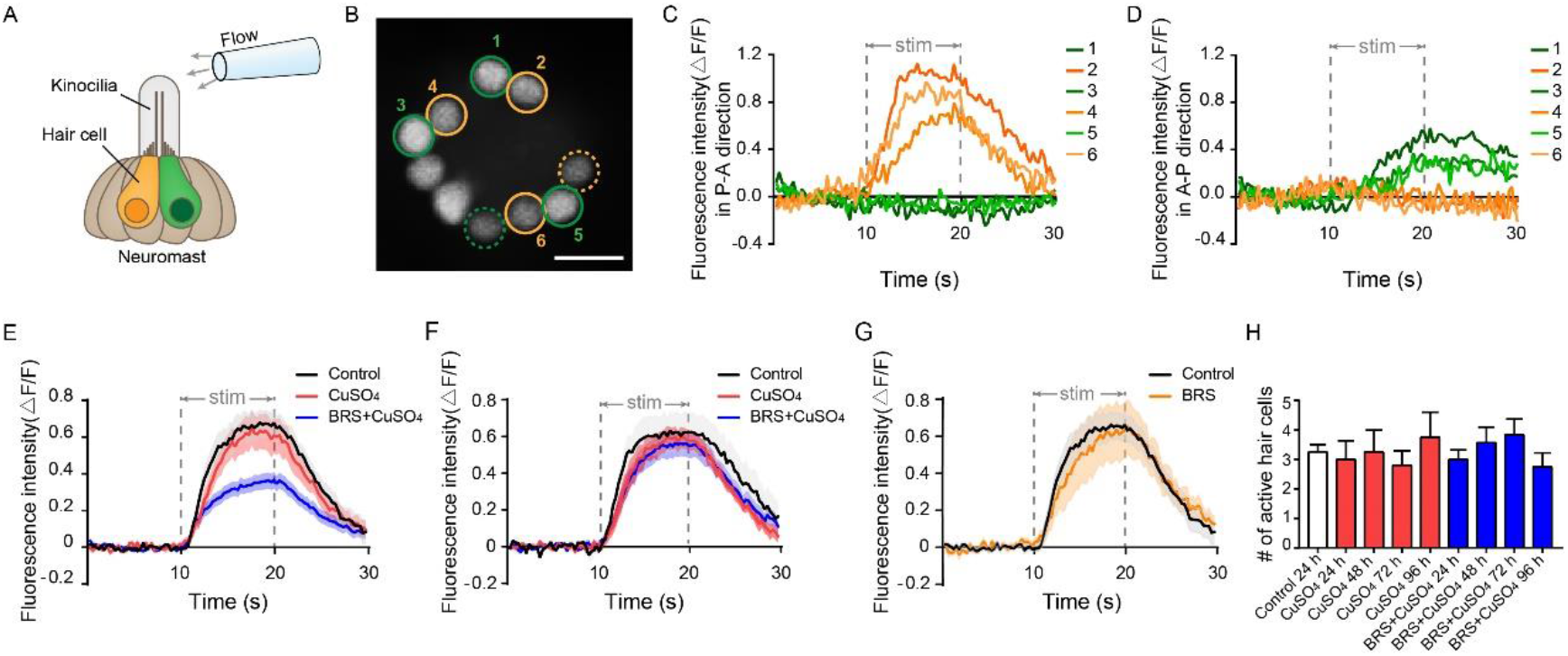
Calcium imaging revealed the function of a single neuromast after hair cell regeneration. **(A)** Schematic diagram shows an electrode filled with fluid is located about 100 μm away from the top of kinocilia to stimulate the neuromast. The yellow and green hair cells represent different polarities. **(B)** When stimulated by the flow, only a part of hair cells respond in this focal plane (circled cells), and some are far from this focal plane (dashed circled cells). The No. 2, 4, and 6 active hair cells (yellow circles) only respond to the flow in P-A direction **(C)**. At the same time, the No. 1, 3, and 5 active hair cells (green circles) only respond to the flow in A-P direction **(D).** Scale bar in represents 10 μm. **(E)** The fluorescence intensity (ΔF/F) of the BRS + CuSO_4_ group is significantly lower than that of the CuSO_4_ group in the Early Stage of regeneration (within 48 hpi)(P < 0.001). **(F)** The ΔF/F of BRS+CuSO_4_ group is not significantly different from that of Control group and CuSO_4_ group in the Late Stage of regeneration (72-96 hpi) **(G)** There is no difference in ΔF/F between the BRS group and the Control group. **(H)** During the regeneration process, the number of active hair cells in CuSO_4_ and BRS+CuSO_4_ group is basically the same, and did not increase with the total number of regenerated hair cells. To **(E-H)**, comparisons were performed by using one-way ANOVA, with Tukey’s multiple comparisons test.

Similar to the results of the rheotaxis, the fluorescence intensity (ΔF/F) of the regenerated hair cells were reduced significantly when inflammation was suppressed at the Early Stage of regeneration (within 48 hpi)(**Fig. 5E**). It was noteworthy that compared to Control group, the fluorescence intensity in CuSO_4_ group did not decrease significantly in the Early Stage of regeneration. This might explain that why the rheotaxis in CuSO_4_ group had been recovered at 24 hpi. The fluorescence intensity of BRS+CuSO_4_ group was not significantly different from that of Control group and CuSO_4_ group in the Late Stage of regeneration (72-96 hpi) (**Fig. 5F**). Additionally, the fluorescence intensity showed no differences between the BRS group and Control group (**Fig. 5G**), indicating that BRS-28 had no effect on the fluorescence intensity.

Normally, only a part of the hair cells in the neuromast responds to the stimulation of water flow. Is it the same for regenerated hair cells? We found that only a few regenerated hair cells in CuSO_4_ group and BRS+CuSO_4_ group responded to flow stimuli. The number of active cells in each neuromast in these two group were approximately the same at 24 to 96 hpi, and were consistent with that in Control group (**Fig. 5H**).

Furthermore, we noticed that most hair cells that responded to the flow in the opposite direction came in pairs (**Supplementary Fig. 3A**). Although the numbers of hair cells responding to flow in P-A direction were similar to that in A-P direction, the fluorescence intensity of hair cells responding to P-A direction was significantly higher than that of hair cells responding to A-P direction (**Supplementary Fig. 3B**). It indicated that L3 neuromast was more sensitive to the flow from the P-A direction.

The results altogether demonstrated that the recovery of hair cell function was delayed at the Early Stage of regeneration when inflammation was suppressed.

## Discussion

### BRS-28 suppresses inflammation and delays the initiation of hair cell regeneration

Although the downregulation of Notch signal during lateral line regeneration induces the proliferation of support cells by activating Wnt/b-Catenin signal (Romero-Carvajal et al., 2015), it is still unknown how the downregulation of Notch signal is triggered after hair cell death. Kniss et al. propos a hypothesis of triggering hair cell regeneration (Kniss et al., 2016). Studies in Drosophila wing disc and eye have found that JNK, Shh, EGF, and TNF signaling pathways are required during apoptosis-induced compensatory proliferation (Fan et al., 2014; Perez-Garijo et al., 2009; Ryoo et al., 2004). Kniss et al. assume that a similar process may be involved in the regeneration of hair cells. On the basis of this hypothesis, we speculate that when hair cells are damaged by CuSO_4_, it cause apoptosis in lateral line hair cells, trigger the rise of reactive oxygen species (ROS) and reactive nitrogen species (RNS), and induce the oxidative stress. This process may improve AP-1,HIF-1α and NF-κB activity, and thus increase pro-inflammatory cytokines and chemokines, such as NO, IL-1β, TNF-α, cox-2,iNOS and so on (Pereira et al., 2016). BRS-28, suppress the production of NO, IL-1β, TNF-α, cox-2, iNOS (Yang et al., 2014), reducing the number of neutrophils and macrophages migrating to the damage of neuromasts. Besides that, the decrease of pro-inflammatory factors may reduce the activation of macrophages. These processes would decrease the production of TNF ligands and inhibit the JNK signal, which contributes to initiating cells regeneration, and eventually leads to delay initiation of compensatory proliferation and delay regeneration of hair cells.

We found that when the initiate time of inflammatory inhibitors was changed, there was no delay in hair cell regeneration (**Fig. 3E**). This also suggests that the timing of inflammation suppression is important: when inflammation occurs, compensatory proliferation of the support cells is triggered and hair cells begin to regenerate. If inflammation suppression does not take effect, regeneration seems to be unaffected.

In addition, neutrophils can also remove dead cell debris, and macrophages can phagocyte apoptotic neutrophils or fragments of dead cells. We believe that when the number and activity of neutrophils and macrophages decrease, the clearance of damaged tissue areas slows down, and hair cells cannot obtain a good regeneration environment. Since damaged neuromasts need more time to clean up these cell fragments, this may also delay the regeneration of hair cells.

### Suppression of inflammation delays functional recovery of regenerated hair cells

In this study, we found that when inflammation was suppressed, hair cell regeneration was delayed, as was the recovery of function. Finally, the quantity and the function of hair cells returned to normal level at the Late Stage of regeneration. Therefore, although the suppression of inflammation delayed the regeneration of hair cells, it did not affect the overall process of hair cell regeneration, and the function of regenerated hair cells eventually tended to be intact. However, the effect of inflammation on the regeneration of lateral hair cells seems to be different from that of the fin. Li et al. found that when zebrafish larvae lack macrophages, vacuoles appear in the regenerated fin, suggesting that macrophages may also be involved in fin regeneration (Li et al., 2012). In our research, although the suppression of inflammation delay regeneration of hair cells and their functional recovery at the Early Stage of regeneration, they eventually return to the normal status at the Late Stage of regeneration. This is not because inflammation is not suppressed sufficiently, as Carrillo et al. found that the number of hair cells finally completed regeneration even when macrophages is knockout (Carrillo et al., 2016). However, this may be because the injured organs are different, and the intact function of lateral hair cells is crucial for the survival of zebrafish. It is suggested that the hair cells in lateral line may have more complex regulation mechanisms during the regeneration process.

### The functional recovery of hair cells is much faster than its quantity

Previous studies have focused on the morphological and quantitative recovery of regenerated hair cells in zebrafish (Carrillo et al., 2016; Romero-Carvajal et al., 2015). Since the regeneration takes 3-4 days post injured, it is easy to assume that the recovery of the function of the neuromasts may be proportional to the number of regenerated hair cells. In this study, for the first time, we performed a method to evaluate the function of regenerated hair cells. We found that the CuSO_4_ group already performed excellent rheotaxis at 24 hpi (**Fig. 4C**), even though the average number of hair cells was only 3.667 at that time (**Fig. 3B**). Therefore, hair cells recover the function much more quickly than their numbers. In other words, although it takes 72-96 h to complete regeneration, the function of hair cells can be recovered within 24 hours which is critical for the survival of zebrafish. When BRS-28 is used to suppress the inflammation, the amplitude of calcium activity of hair cells is significantly lower than that of Control and CuSO_4_ group at the Early Stage of regeneration, and the rheotaxis of larvae is poor during this period. Therefore, the suppression of inflammation not only delays the hair cell regeneration, but also delays the functional recovery.

There is a mismatch between the function and quantity during regeneration. Calcium image reveal that only a part of regenerated hair cells in one neuromast respond to the flow. This finding is consistent with previous study (Zhang et al., 2018). In our research, we found that this phenomenon also exists in regeneration group (CuSO_4_ and BRS+CuSO_4_ group). Regardless of the number of regenerated hair cells, the number of hair cells that respond to water flow remain stable during the regeneration process, which is not different from Control group (**Fig. 5H**). Besides that, in the Early Stage of regeneration, the magnitude of fluorescence intensity and reaction time of CuSO_4_ group are also consistent with that of the controls. This explains why the number of regeneration in the CuSO_4_ group at 24 h is only 3.667 on average, but the function of the lateral line has been restored to a level very close to that of Control group.

In this study, we only performed calcium imaging on the L3 neuromast, which was confirmed as the polarity of the A-P body axis in the study of Chou et al (Chou et al., 2017). Consistent with their results, this neuromast is indeed insensitive to the flow in the dorsal-ventral (D-V) body axis (data not shown). Therefore, this study only focuses on the stimulus response in the A-P body axis direction, and does not further analyze the stimulus data in the D-V body axis direction. Compared with hair cells with polarity in the A-P direction, hair cells with polarity in the P-A direction have greater ΔF/F0 when stimulated by water flow (**Supplementary Fig. 3B**; sample, **Fig. 5 C, D**). It indicated that L3 neuromast is more sensitive to the flow from the P-A direction. This finding is consistent with the results measured by Chou et al. using microphonic potentials evoked by sinusoidal stimuli (Chou et al., 2017). Most active hair cells that responded to the opposite flow come in pairs (**Supplementary Fig. 3A**), suggesting that it appears to be pre-arranged rather than random.

In summary, our research suggests that suppression of inflammation delays functional regeneration of lateral hair cells in zebrafish larvae. The inflammation plays positive and permissive roles in hair cell regeneration.

## Materials and Methods

### Zebrafish strains and maintenance

AB/Wild-type strain, Tg(corola-eGFP;lyz-Dsred) and Huc:h2b-gcamp6f transgenic line were used in this study. Embryos were generated by paired mating and maintained at 28.5°C in EM and on a 14/10 h light/dark cycle according to the standard protocols.

All animal manipulations were conducted strictly in accordance with the guidelines and regulations set forth by the University of Science and Technology of China (USTC) Animal Resources Center and the University Animal Care and Use Committee. The protocol was approved by the Committee on the Ethics of Animal Experiments of the USTC (Permit Number: USTCACUC1103013).

### Hair cell damage and inflammation inhibition

In order to damage hair cells in lateral line, 4 dpf Larvae were treated with 5 μM CuSO_4_ (Sangon, China) diluted in embro medium (EM) for 1 h. Then, they were washed three times and recovered in EM.

To suppress inflammation, 4 dpf larvae were immersed in 0.1% BRS-28, an anti-inflammatory agent, for 3 h before being moved into CuSO_4_ to damage hair cells.

### Live imaging

AB/Wild-type larvae were used to count the number of regenerated hair cells in L2, LII3, L3 neuromasts (**Fig. 1A**). Hair cells were marked by 0.01 %DAPI for 5 minutes. Larvae were anesthetized in 0.02% MS-222 (Tricaine mesylate, Sigma, USA) and imaged under a fluorescence microscope (Olympus BX-60, Japan).

In order to exhibit the damage of hair cells in copper sulfate solution and the regeneration of hair cells in different phases, hair cells were labeled by 0.05 % DASPEI (Sigma, USA), and larvae were anesthetized in MS-222 and imaged under a confocal microscopy (Zeiss LSM 880 +Airyscan, Germany).

Tg(corola-eGFP; lyz-Dsred) transgenic line was used to observe the number of neutrophils and macrophages migrating to the injured neuromasts *in vivo*. In this transgenic line, neutrophils co-expressed *lyz-Dsred* and *coro1a-GFP* and show yellow fluorescence after these two channels are merged, while macrophages only express *coro1a-GFP* and show green fluorescence (Li et al., 2012). To show the neutrophils and macrophages migrating to damaged neuromasts, larvae were anesthetized in MS-222 and imaged under a confocal microscopy (Zeiss LSM 880 +Airyscan). In order to count neutrophils and macrophages, we set the area around the L2 LII3 L3 neuromasts with a diameter of 100 μm as the region of interest (ROI). Zebrafish larvae were anesthetized and imaged by the fluorescence microscope (Olympus BX-60) with a green and a red channel.

### Rheotaxis behavior experiments

A U-shaped tank was designed to test the rheotaxis behavior of larvae (**Fig. 4A**). The bottom of the two cubic tanks (7 cm length *8 cm width*8 cm height) were connected by a platform (10 cm length *8 cm width*0.5 cm height). A peristaltic pump (Longer Pump YZ1515x, China) was used to move EM solution from the left tank to the right tank, so that the platform formed a steady water flow from right to left (v=10 mm/s). AB/WT zebrafish larvae were applied to detect the ability of rheotaxis. Larvae were released at the right side of the platform with an initial velocity almost equals 0. To avoid visual cues, experiments were operated in the dark and rheotaxis performs were recorded by an infrared CCD (IR850, weixinshijie, China).

Rheotaxis data were analyzed by our own rheotaxis software edited in Matlab (2015a, MathWorks, USA). This software can plot the movement track of zebrafish larvae in the platform, measure the direction and distance of each swimming and calculate the speed. Finally, it reports scores based on the magnitude in the horizontal direction of the ratio of motion vectors sum to the motion arithmetic sum.

### Calcium imaging and data analyses

Huc: h2b-GCamp6f transgenic line was used in calcium imaging which expressed pan-neuronal nucleus-labelled GCamp6f. Larvae were anesthetized and fixed by a net pressure. The one-step pulled micropipette had a long, wispy tip which must be trimmed by rubbing it against with another pulled micropipette to generate a tip with an outer diameter of approximately 40 μm. The micropipette was filled with 0.02% MS-222 and fixed to the holder of a micromanipulator (MX7500, Scientific Design Company, USA). The tip of the micropipette was positioned at a distance of approximately 100 μm from the top of the kinocilia (**Fig. 5A**). The duration of flow was controlled by three direct links which were linked with a syringe.

Calcium imaging was collected by a confocal microscopy (FV 1000, Olympus, Japan). To make as many hair cells as possible in the observation area at the same time, a single z-axis was adjusted. ROI was set to 110*108. We took 100 time-lapse images for each neuromast, and the total capture time was 29.7 s (0.297 s per slice). Flow stimulation occurred from 10.098 to 19.899 s.

Since the neuromasts are three-dimensional, different hair cells have different levels of fluorescence intensity. Namely, they have different levels of F prime. The relative fluorescence intensity change (ΔF/F_0_) is more commonly used. For each hair cell, the average fluorescence intensity before flow stimuli (0-10 s) was set as F_0_. The data would be excluded when F_0_<95, which means these hair cells were too far from the focal plane. When more than two hair cells in the neuromast respond to flow stimulation, two hair cells with the strongest fluorescence were selected and included in the statistics of fluorescence intensity curve.

### Statistical analysis

All data were shown as mean ± S.E.M. or as relative proportions of 100 % as indicated in the appropriate legends. The data were analyzed in either one-way ANOVA with Tukey’s multiple comparisons test or two-way ANOVA with Tukey’s multiple comparisons test by GraphPad Prism version 7.0 (Prism, San Diego, CA, USA). The level of significance was set to P < 0.05. *, **and ***represent P < 0.05, P < 0.01 and P < 0.001, respectively.

## Acknowledgments

The authors thank Drs. Wen Zilong for providing the Tg(corola-eGFP; lyz-Dsred) transgenic fish line, Drs. Wen Quan for providing the Huc:h2b-gcamp6f transgenic fish line. The authors thank Drs. Zhen Xuechu for providing BRS-28 and the compound-26 in their study is the BRS-28 mentioned in this study.

---

**Supplementary Fig. 1.**
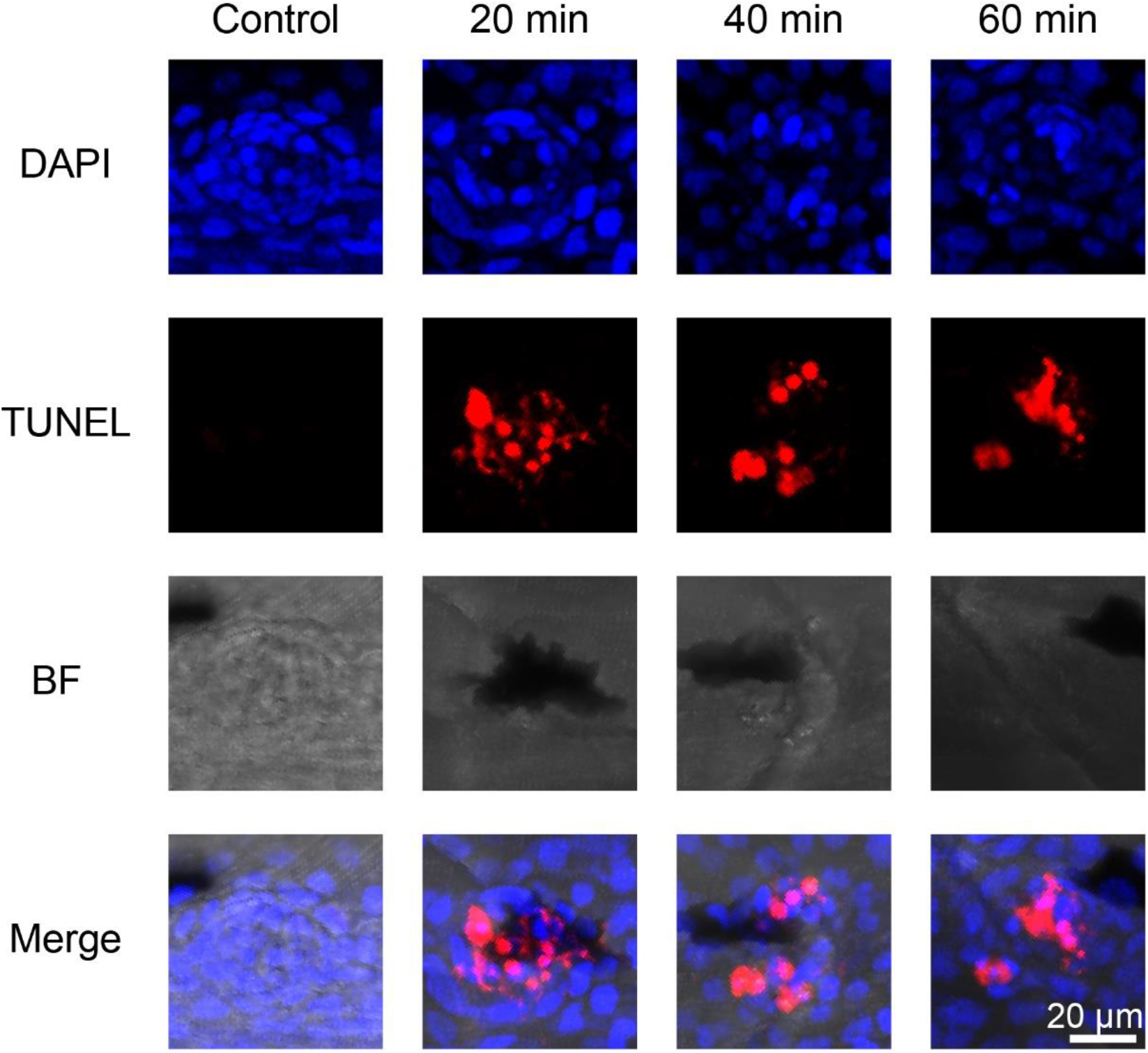
CuSO_4_ caused apoptosis in hair cells. TUNEL assay revealed hair cells occurred apoptosis when treated with CuSO_4_. Nuclei were stained with DAPI. BF: Bright Field. Scale bar represents 20 μm.

**Supplementary Fig. 2.**
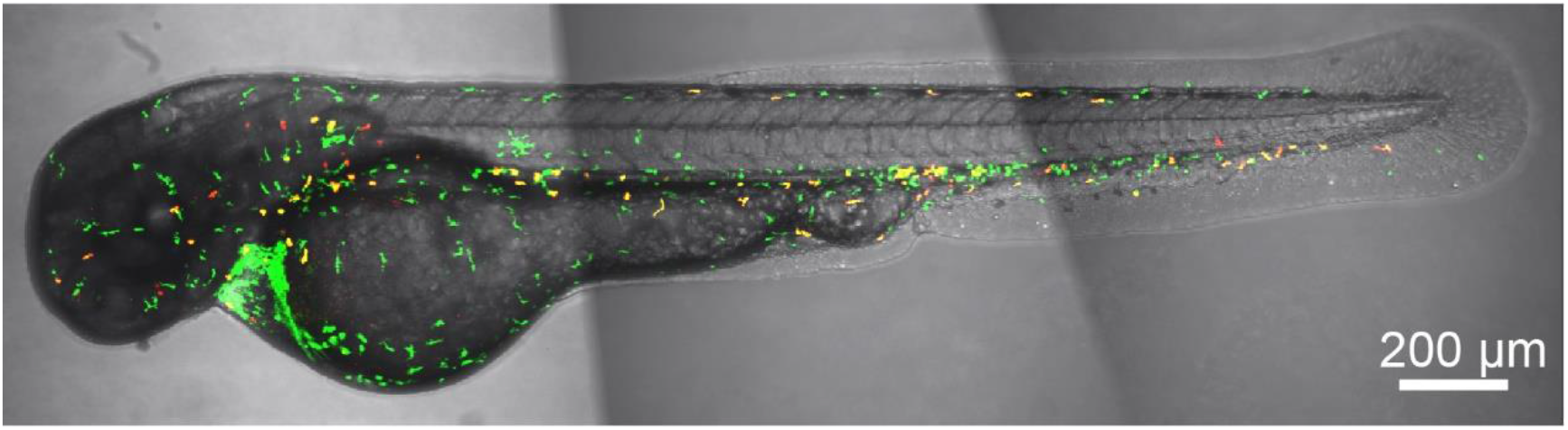
Tg(corola-eGFP; lyz-Dsred) transgenic line could mark both neutrophils and macrophages. In Tg(corola-eGFP; lyz-Dsred) transgenic line, neutrophils co-expressed *lyz-Dsred* and *coro1a-GFP* and show yellow fluorescence after these two channels are merged, while macrophages only express *coro1a-GFP* and show green fluorescence. Scale bar represents 200 μm.

**Supplementary Fig. 3.**
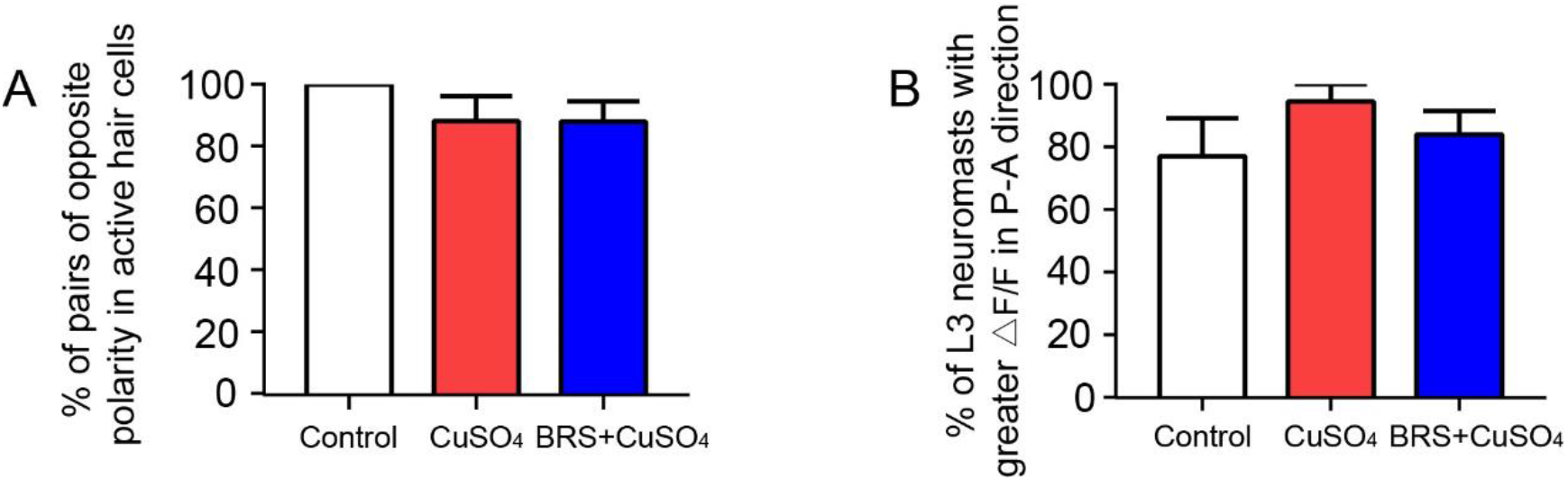
Most active hair cells are polar in pairs and are sensitive to flow in the P-A direction. **(A)** Most hair cells that responded to the flow in the opposite direction come in pairs. **(B)** The fluorescence intensity of hair cells responding to P-A direction is significantly higher than that of hair cells responding to A-P direction. To **(A,B)**, comparisons were performed by using one-way ANOVA, with Tukey’s multiple comparisons test. All Error bars show mean ± S.E.M.

